# Targeting the Cx26/NANOG/Focal Adhesion Kinase Complex via Cell-Penetrating Peptides in Triple-Negative Breast Cancers

**DOI:** 10.1101/2021.09.09.459689

**Authors:** Erin E. Mulkearns-Hubert, Emily Esakov Rhoades, Salma Ben-Salem, Rashmi Bharti, Nicole Hajdari, Sarah Johnson, Alex Myers, Iris Nira Smith, Smarajit Bandyopadhyay, Charis Eng, Erinn Downs-Kelly, Justin D. Lathia, Ofer Reizes

## Abstract

Triple-negative breast cancer (TNBC) represents the most lethal and treatment-resistant breast cancer subtype with limited treatment options. We previously identified a protein complex unique to TNBC cancer stem cells composed of the gap junction protein connexin 26 (Cx26), the pluripotency transcription factor NANOG, and focal adhesion kinase (FAK). We sought to determine whether a peptide mimetic of Cx26 designed to target the complex attenuated tumor growth in pre-clinical models. Histological assessment was employed to verify expression of complex members. We designed peptides based on Cx26 juxtamembrane domains and performed binding experiments with NANOG and FAK using surface plasmon resonance. Peptides with high affinity were engineered with a cell-penetrating sequence and assessed in functional assays including cell proliferation, self-renewal, and in vivo tumor growth, and downstream signaling changes were measured. Binding studies revealed that the Cx26 C-terminal tail and intracellular loop bound to NANOG and FAK with submicromolar-to-micromolar affinity and that a 5-amino acid sequence in the C-terminal tail of Cx26 (RYCSG) was sufficient for binding. The Cx26 C-terminal tail was tagged with an antennapedia cell-penetrating peptide sequence and intracellular localization was confirmed. The cell-penetrating Cx26 peptide (aCx26-pep) disrupted self-renewal as assessed by tumorsphere formation assay while reducing nuclear FAK and NANOG and inhibiting NANOG target gene expression in TNBC cells but not luminal mammary epithelial cells. In vivo, aCx26-pep reduced tumor growth and proliferation and induced cell death. We provide proof-of-concept that a Cx26 peptide-based strategy inhibits growth and alters NANOG activity in TNBC.

## Introduction

Triple-negative breast cancer (TNBC), the most aggressive breast cancer subtype, is associated with high rates of recurrence and metastasis^1-5^. To date, unlike other breast cancer subtypes, no specific targeted approaches exist for TNBC, and toxic chemotherapeutic agents are the primary treatment regimen, highlighting the need to develop new targeted therapies. TNBC contains a high degree of cellular heterogeneity and plasticity that has emerged as a hallmark of the malignant state and accounts for the observed persistent tumor growth, therapeutic resistance, and metastasis^6-9^. TNBC also contains self-renewing, therapeutically resistant cancer stem cells (CSCs) that are responsible for tumor progression and metastasis^10-13^. Given the limited treatment options for TNBC, CSC neutralizing strategies are an immediate priority for therapeutic development.

Attempts to elucidate the complexity of CSC maintenance have focused on intrinsic driver mutations and altered developmental signaling pathways^14^. The elevated cellular density within tumors stimulates cellular programs that are activated by cell-cell contact and close proximity. The gap junction family of proteins is composed of connexin subunits and canonically functions in gap junction plaques at the interface of adjacent cells to facilitate direct cell-cell communication. Connexins can also function non-canonically as single membrane channels (hemichannels, which contain a single assembled connexon hexamer of connexins) or as signaling hubs adjacent to any organelle and/or the plasma membrane^15^. In cancer, the classical role of connexins as tumor suppressors has been widely described in many tumor models^16-20^. However, the paradigm that connexins have a global tumor-suppressive role has been challenged by recent evidence that connexins can be pro-tumorigenic by facilitating tumor progression and metastasis^21-27^. The connexin subunits involved in maintaining homeostasis during the physiological development and function of the mammary gland are Cx26 and Cx43^18, 28^. In breast cancer, connexins have been described to be both pro- and anti-tumorigenic by regulating transformation, proliferation, cell survival, and metastasis^27^. Most studies to date suggest a tumor-suppressive role for Cx26 early in breast cancer progression based on evidence that Cx26 is frequently absent or down-regulated in human breast cancer cell lines and human primary tumors^15, 29-31^. However, these studies are inconsistent with clinical observations that demonstrate a strong correlation between increased Cx26 expression in breast cancer tissue samples post-treatment (chemotherapy/surgery) and decreased overall survival^32^. Moreover, Cx26 expression has been shown to be associated with increased lymphatic vessel invasion, increased tumor size, and poor prognosis in human breast cancers^32, 33^.

We previously demonstrated that Cx26 is elevated in TNBC compared to normal mammary tissue and enriched in CSCs compared to their non-CSC progeny in TNBC cell lines and patient-derived xenograft models^34^. In functional studies, we demonstrated that Cx26 is necessary and sufficient for CSC maintenance and regulates NANOG protein stability. In TNBC, Cx26 localizes to an intracellular membrane-bound vesicle in complex with the pluripotency transcription factor NANOG and focal adhesion kinase (FAK). In contrast to its role in the normal mammary epithelium, Cx26 functions in a gap junction-independent manner in TNBC by forming an intracellular complex that underlies the maintenance of CSC function and stabilizes NANOG. We leveraged these initial findings to develop a peptide-based strategy to disrupt this novel signaling network. We developed peptides to the intracellular and extracellular regions of Cx26 and identified the binding domains necessary for interaction with NANOG and FAK. Using a cell-penetrating peptide strategy, we show that introduction of the Cx26 C-terminus into TNBC cells inhibits self-renewal, reduces NANOG target gene expression, and attenuates in vivo tumor growth. Together, this study provides proof-of-concept that targeting CSCs via connexin peptide mimetics represents a viable therapeutic strategy for TNBC.

## Materials and Methods

### Study Approvals

All mouse procedures were performed under adherence to protocols approved by the Institute Animal Care and Use Committee at the Lerner Research Institute, Cleveland Clinic (#2018-2101). All human specimens were obtained under adherence to protocols approved by the Cleveland Clinic Foundation Institutional Review Board (IRB approval #17-418).

### TNBC Cx26, FAK, and NANOG Histology and Analysis

Annotated cases of invasive ductal carcinoma were retrieved from the surgical pathology files of Cleveland Clinic and included triple-negative breast carcinoma (TNBC), hormone receptor positive (HR+ being estrogen receptor positive) and HER2 negative (HR+/HER2-), hormone receptor negative (HR-being estrogen receptor negative) and HER2 positive (HR-/HER2+) or hormone receptor positive and HER2 positive (HR+/HER2+). In total, 12 TNBC, 12 HR+/HER2-, 5 HR-/HER2+, 4 HR+/HER2+ tumors were evaluated. Hematoxylin and eosin stained slides were reviewed for invasive tumor adequacy and immunohistochemistry (IHC) was performed on 4-μm thick formalin-fixed paraffin-embedded (FFPE) whole tissue serial sections following antigen retrieval (Target Retrieval Solution, pH 6.1; Dako, Carpinteria, CA). Immunostains performed included Connexin 26 (Cx26) mouse monoclonal antibody (clone CX-12H10, Thermo Fisher Scientific, Waltham, MA) at 1:200 dilution, NANOG rabbit monoclonal antibody (clone EPR2027, Abcam, Cambridge, MA) at 1:250 dilution, and FAK rabbit polyclonal antibody (clone Q05397, Cell Signaling Technology, Danvers, MA) at 1:500 dilution. All staining was performed on an automated platform (Ventana Benchmark, Ultraview XT, Ventana Medical Systems, Inc, Tucson, AZ) and visualized with ultraView universal DAB detection kit (Ventana Medical Systems, Inc, Tucson, AZ). The cellular compartment staining (membranous, cytoplasmic or nuclear) was evaluated and the intensity in each case for the three antibodies was characterized as weak, moderate, or strong, and the extent of immunoreactivity scored according to the percentage of tumor cells with staining: 0 (0%), 1+ (<5%), 2+ (5–25%), 3+ (26–50%), 4+ (51–75%), and 5+ (76–100%).

### Cell Culture

MDAMB231, MDA-MB-468, HCC70, and MCF7 breast cancer cells (American Type Culture Collection; Manassas, VA) were cultured in log-growth phase in Dulbecco’s modified Eagle’s medium (DMEM) supplemented with 1 mM sodium pyruvate (Cellgro, Kansas City, MO) and 10% heat-inactivated fetal calf serum (FCS) at 37°C in a humidified atmosphere (5% CO_2_). All cell lines were authenticated through the Cleveland Clinic Lerner Research Institute Cell Culture Core and routinely tested for mycoplasma contamination. The NANOG-GFP reporter was described previously, and cells were sorted for GFP^hi^ and GFP^lo^ populations as described^35^.

### Peptide Synthesis and Validation

All peptides, unless otherwise noted, were synthesized by the LRI Molecular Biotechnology Core through standard solid-phase chemistry, using Rink Amide resin and side chain-protected Fmoc-amino acids, on a Liberty Blue Automated Microwave Peptide Synthesizer (CEM Corporation, Matthews, NC). Each peptide was purified□to >□95% by acetonitrile gradient elution (20 - 50%) on a Vydac preparative C18 column, using reverse-phase HPLC (Waters Corporation). The quality of the peptide was analyzed by HPLC (Agilent 1100) on a reverse-phase analytical C-18 column. The correct mass corresponding to the amino acid composition of each peptide was then confirmed by mass spectrometry (MALDI-TOF/TOF mass spectrometer; Model No. 4800; AB SCIEX, Framingham, MA). The peptides were lyophilized and stored at -80°C until reconstituted in 1% DMSO/PBS.

### 3D protein structure prediction

AlphaFold2 was used to predict the 3D structure of Cx26, NANOG and FAK: (https://www.blopig.com/blog/2021/07/alphafold-2-is-here-whats-behind-the-structure-prediction-miracle/). Images were rendered using Visual Molecular Dynamics software^36^.

### Surface Plasmon Resonance (SPR) Analysis

The equilibrium dissociation constants (KD) between synthesized Cx26 cytosolic tail peptides and either NANOG or FAK proteins were determined using the Biacore S200 (Cytiva, USA) at 25°C. The 1% DMSO/1x PBS-P+ (Cytiva) buffer was filtered through a 0.22-μm membrane filter and degassed immediately before the experiment as running buffer with a flow rate of 30 μL/min for the instrument system. The purified active NANOG was diluted in 10 mM sodium acetate solution pH 4.5, resulting in a protein concentration of 50 μg/mL, and purified active FAK was diluted in 10 mM sodium acetate solution pH 5.0, resulting in a protein concentration of 20 μg/mL. The diluted protein was immobilized on the surface of a Series S Sensor Chip CM5 via the primary amine group, employing a standard Amine Coupling Kit (1-Ethyl-3-(3-dimethylaminopropyl)- carbodiimide hydrochloride (EDC), N-hydroxysuccinimide (NHS) and 1.0 M ethanolamine-HCl (pH 8.5), and the target immobilization level was 5000 response units (RUs). The flow cell 1 without any modification was used as a reference to correct for system error. The RU values were collected, and all the binding affinity data were calculated by kinetic models (1:1 Langmuir interaction) within the Biacore S200 Evaluation Software. To determine the binding affinity between each peptide and NANOG or FAK, a series of peptide dilutions was analyzed by multi-cycle kinetics/affinity. A concentration gradient of each peptide as the analyte was freshly prepared in running buffer over at least five concentrations (100 nM – 100 μM). Each peptide at various gradient concentrations (including a repeat concentration) and one zero concentration (running buffer) flowed over the immobilized NANOG or FAK, with 180 s for binding, followed by disassociation for 600 s, and the obtained RUs were recorded. After the injection of each concentration of each peptide, the sensor chip surface was regenerated with 10 mM glycine pH 2.5 for 30 s to completely remove the residual peptide. The KD was calculated to evaluate the ability of peptides to interact with NANOG or FAK proteins.

### Immunofluorescence and Flow Cytometry Analysis of Antp-peptides

Following peptide treatment, MDA-MB-231 cells were lifted and washed in 1x PBS followed by staining with UV Live/Dead dye (1:1000, Thermo Scientific) for 15 min at room temperature. Cells were then fixed and permeabilized using the Annexin V Staining Kit (BD Biosciences) per the manufacturer’s instructions and incubated with FITC-conjugated anti-antennapedia (anti-Antp-FITC) antibody (1:100, LSBio) for 30 min. Samples were washed and analyzed using a BD Fortessa flow cytometer. Data analysis was performed using the FlowJo software v10 (Tree Star, Inc., Ashland, OR).

For immunofluorescence staining, MDA-MB-231 cells were plated onto glass coverslips coated with Geltrex (ThermoFisher) in 24-well plates and treated with aCx26-pep (10 μM) for 16 h. Subsequently, coverslips were washed with 1x PBS, fixed with 4% paraformaldehyde, and either permeabilized with 0.1% Triton X-100 or left non-permeabilized. Cells were then stained with anti-antennapedia antibody (anti-Antp 4C3, 1:100, University of Iowa Developmental Studies Hybridoma Bank) for 1 h at room temperature. Coverslips were washed twice with 1x PBS and stained with Texas Red-conjugated anti-mouse secondary (1:1000, Invitrogen) for 1 h at room temperature in the dark, before being mounted in DAPI mount (VWR) onto glass slides and imaged with a confocal microscope at 63x magnification (Leica). Appropriate secondary alone controls were completed to confirm the absence of background staining.

### CellTiter Glo Proliferation Studies

MDA-MB-231 cells were plated in a 96-well (opaque-walled) plates at 1,000 cells/well and treated for 72 h with Antp peptide alone, aCx26-pep or vehicle (1% DMSO in PBS) at the concentrations shown in the figures. At day 0 and post 72 h of treatment, cells were lysed with CellTiter-Glo® Luminescent Cell Viability Assay reagent (Promega), shaken in the dark for 10 min, and luminescence was recorded. Fold change was calculated by normalizing the 72 h luminescence value to day 0 for each group.

### Self-renewal (Tumorsphere) Assays

Duplicate rows of cells were cultured in serial dilutions per well (1-20 cells per well) in one 96 well plate (Sarstedt, Germany) per condition with 200 μl serum-free DMEM supplemented with 20 ng/ml basic fibroblast growth factor (Invitrogen, Grand Island, NY), 10 □ng/ml epidermal growth factor (BioSource, Grand Island, NY), 2% B27 (Invitrogen, Grand Island, NY), and 10 μg/ml insulin. The frequency of sphere formation was calculated in such a way that a well with a tumorsphere was counted as a positive well and a well with no tumorspheres was counted as a negative well. Tumorspheres were counted under a phase-contrast microscope after 2 weeks. The stem cell frequencies were calculated using an extreme limiting dilution algorithm (http://bioinf.wehi.edu.au/software/elda/)^37^.

### Subcellular Fractionation and Immunoblotting

A total of 15×10^6^ cells were plated in a 15 cm dish and treated with 10 μM peptide or 1% DMSO vehicle for 48 h. Cells were subsequently lysed and subcellular fractions prepared using the Subcellular Fractionation Kit for Cultured Cells (ThermoFisher) according to the manufacturer’s protocol. A total of 25 μg of each fraction was loaded on a 12% SDS-PAGE gel and transferred to PVDF membrane. Membranes were blocked in 5% non-fat milk in TBST for 1 h and incubated with the appropriate primary antibody overnight at 4°C: anti-NANOG (D73G4) (Cell Signaling #4903; 1:1000), anti-FAK (D2R2E) (Cell Signaling #13009; 1:1000), anti-Lamin A/C (Proteintech #10298-1-AP; 1:3000). Membranes were then washed in TBST and incubated with the appropriate HRP-tagged secondary antibody (Sigma-Aldrich) for 3 h at room temperature. Rhodamine-tagged anti-actin (Bio-Rad #12004163; 1:10,000) was used as a loading control. All immunoblots were imaged on a Bio-Rad ChemiDoc MP.

### Real-time Quantitative PCR

Quantitative polymerase chain reaction (qPCR) was performed using a StepOnePlus™ Real-Time PCR System (Applied Bioystems) with SYBR-Green MasterMix (SA Biosciences). RNA was extracted using the RNeasy kit (Qiagen, Limburg, Netherlands), and cDNA was synthesized using the Superscript III kit (Invitrogen, Grand Island, NY). For qPCR analysis, the threshold cycle (CT) values for each gene were normalized to expression levels of GAPDH. Dissociation curves were evaluated for primer fidelity. The primers used are detailed in Supplemental Table S3.

### Animal Studies

*NOD.Cg-Prkdc*^*scid*^ *Il2rg*^*tm1Wjl*^*/SzJ* (NSG) female mice were purchased from Jackson Laboratories or the Biological Resource Unit at the Lerner Research Institute, Cleveland Clinic. All mice were maintained in microisolator units and provided free access to food and water. MDA-MB-231 cells were transduced with a luciferase lentiviral vector (pHIV-Luciferase, Addgene #21375). Following puromycin drug selection, 1×10^6^ cells were injected into the right subcutaneous flank of 6-week-old female mice. Once tumors became palpable, mice were randomized into groups and received a total of 50 μl of either 0.1 mM peptide or vehicle (1% DMSO) via intra-tumoral injection every other day for 20 days. Tumor progression was assessed weekly via bioluminescent IVIS imaging (see below). Following treatment regiment, mice were euthanized by CO_2_ asphyxiation with subsequent cervical dislocation and necropsied for further analysis.

### Bioluminescent IVIS Imaging

Bioluminescence images were taken with an IVIS Lumina (PerkinElmer) using D-luciferin as previously described^38^. Mice received an intraperitoneal (IP) injection of D-luciferin (Goldbio LUCK-1G, 150 mg/kg in 150 μL) under inhaled isoflurane anesthesia. Images were analyzed (Living Image Software), and total flux were reported in photons/second/cm^2^/steradian for each mouse.

### Immunohistochemistry

Upon necropsy, subcutaneous tumors were excised, sectioned and placed in 4% paraformaldehyde (PFA) for 3 days at room temperature, before being transferred into 70% ethanol and sent to the Histology Core at the LRI for processing, sectioning, and staining. Slides were imaged using the Aperio slide scanner and analyzed.

### Statistical Analysis

Values reported in the results represent mean□±□standard error of the mean (SEM). Oneway analysis of variance was used to calculate statistical significance, and *p* values and replicate numbers are detailed in the text and figure legends. All experiments were repeated at least three separate times, with technical replicates performed within each experimental replicate as appropriate.

### Data Availability Statement

The data generated in this study are available upon request from the corresponding author.

## Results

### The Cx26/NANOG/FAK complex is present in patient TNBC samples

Previous data from our group identified a protein complex unique to TNBC CSCs composed of the gap junction protein connexin 26 (Cx26), the pluripotency transcription factor NANOG, and focal adhesion kinase (FAK)^34^. To assess whether the Cx26/NANOG/FAK complex is also present in tumors from patients, we stained sections of human triple-negative invasive ductal carcinoma for Cx26, NANOG, and FAK. Hematoxylin & eosin staining showed the invasive nature of the tumors (**Fig. 1A**). In cultured TNBC cells, we found that Cx26 localizes to an intracellular compartment rather than the canonical plasma membrane localization of connexins^34^. Consistently, we observed intracellular localization of staining for Cx26, NANOG, and FAK in adjacent tumor sections of TNBC patient tumor (**Fig. 1B-D**). In contrast, tumor samples from patients with hormone receptor-positive and HER2-negative tumors showed cell surface localization of Cx26 distinct from NANOG and FAK (**Fig. 1E-H, summarized in Fig. 1I**). This indicates that Cx26, NANOG, and FAK have similar subcellular localizations in TNBC, suggesting that the complex may be present in TNBC patient tumors as well as in cells, as we previously described^34^.

**Figure 1.**
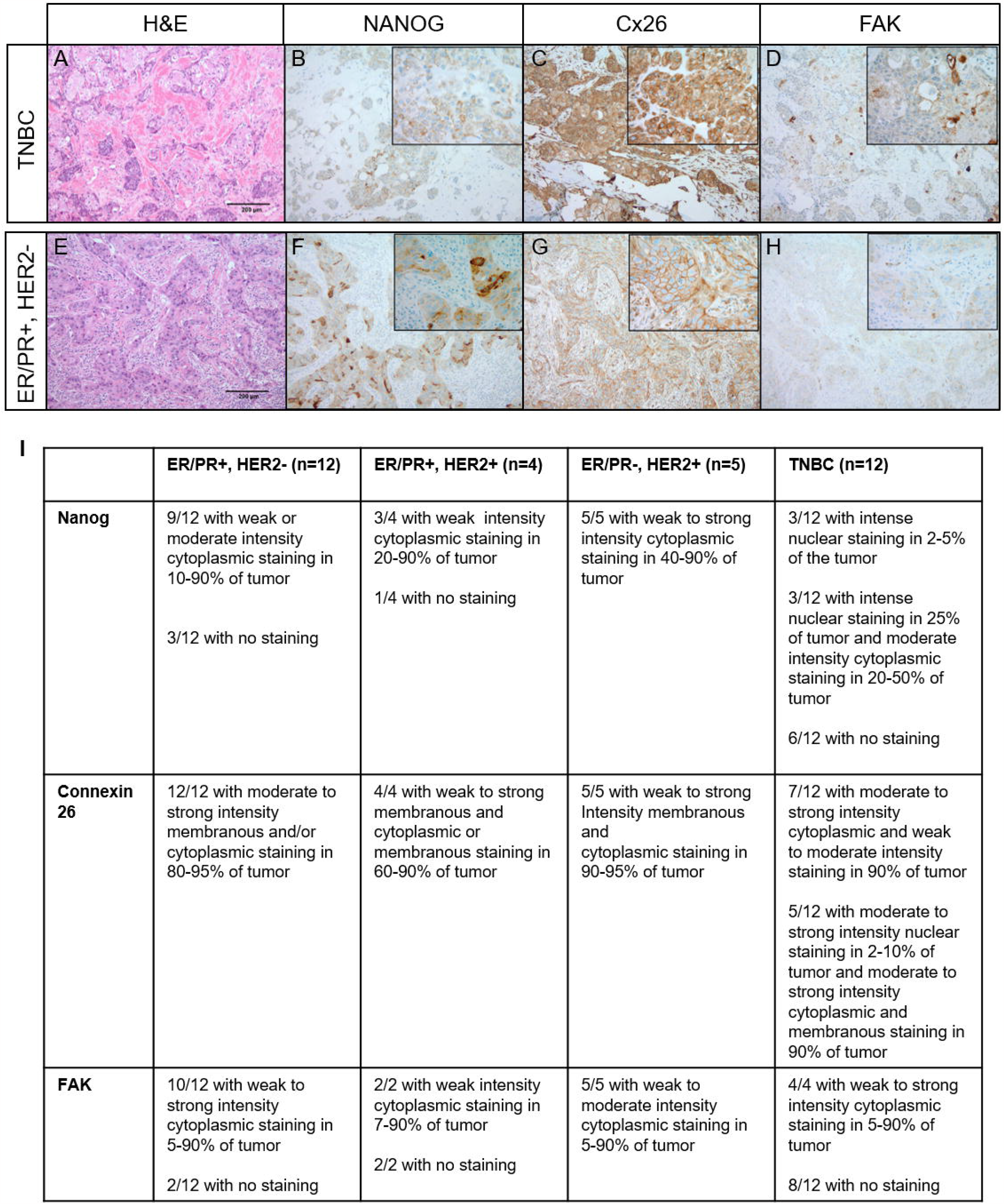
Cx26, NANOG, and FAK localize to intracellular compartments in TNBC tumors. A-D: TNBC assessed by sequential whole slides as stained with A) Hematoxylin and eosin showing an invasive ductal carcinoma with focal squamous differentiation, Nottingham grade 3, B) NANOG stain showing weak cytoplasmic expression within tumor cells (inset showing granular cytoplasmic staining in a large subset of the tumor cells), C) Cx26 stain showing intense cytoplasmic staining within a majority of the tumor cells (inset showing higher magnification) and D) FAK stain showing weak granular cytoplasmic staining within a subset of tumor cells (inset showing weak granular cytoplasmic staining and darker accentuation of staining in area of squamous differentiation). E-H: Estrogen/progesterone receptor (ER/PR)+/HER2-tumor assessed by sequential whole slides stained with E) Hematoxylin and eosin showing an invasive ductal carcinoma, Nottingham grade 3, F) NANOG immunohistochemical stain showing variable cytoplasmic expression within tumor cells (inset showing both weak and more intense foci of cytoplasmic staining), G) Cx26 immunohistochemical stain showing circumferential membranous expression on tumor cells (inset showing moderate to strong diffuse membrane staining) and H) FAK immunohistochemical stain showing very weak granular cytoplasmic staining within tumor cells (inset showing weak but diffuse granular cytoplasmic staining). Photomicrograph total magnification for each pane is 100x; inset total magnification is 400x. I) Summary table describing histological findings.

### FAK and NANOG interact with the cytosolic tail of Cx26

Based on the essential role for the Cx26/NANOG/FAK complex in TNBC CSC maintenance^34^, targeting the complex represents a therapeutic opportunity. To develop therapies targeting the complex, we used surface plasmon resonance (SPR) to interrogate the specific domains through which the proteins interact. We generated peptide fragments representing the intracellular and extracellular domains of Cx26 and determined their ability to bind to either FAK or NANOG by SPR. The strongest binding occurred between NANOG and the Cx26 cytosolic tail and between FAK and both the cytosolic tail and the cytosolic loop, with dissociation constants in the nanomolar and micromolar ranges, respectively (**Fig. 2A**). Importantly, no binding was observed between the FAK and NANOG proteins alone (**Fig. 2A**). These observations combined with predictive modeling suggest that complex binding requires Cx26, as we detected binding between Cx26 and NANOG, Cx26 and FAK, and all members of the complex but not between FAK and NANOG alone (**Supplemental Fig. 1**). These data informed a binding schematic for the Cx26/NANOG/FAK complex (**Fig. 2B**) where NANOG binds to the C-terminus while FAK interacts with both the C-terminus and the intracellular loop of Cx26. To determine the specific Cx26 cytosolic tail amino acid residues important for binding, several truncations of the C-terminal tail were synthesized and subjected to SPR analysis for binding to both NANOG and FAK. The minimal required region mapped to Cx26 amino acids 216-220 (**Supplemental Table S1 and Supplemental Fig. S2 and S3**). To gain more insight into the kinetics of binding, we co-incubated the Cx26 cytosolic tail with NANOG prior to applying the components to a FAK-bound chip. We also performed this analysis with Cx26 cytosolic tail pre-incubated with FAK prior to applying to a NANOG-bound chip. Interestingly, the Cx26 cytosolic tail had decreased binding affinity for both FAK and NANOG when pre-incubated with the other complex member compared to the Cx26 cytosolic tail alone or pre-incubation with bovine serum albumin control (not shown). Taken together, these data indicate that NANOG and FAK binding to the Cx26 cytosolic tail may be competitive.

**Figure 2.**
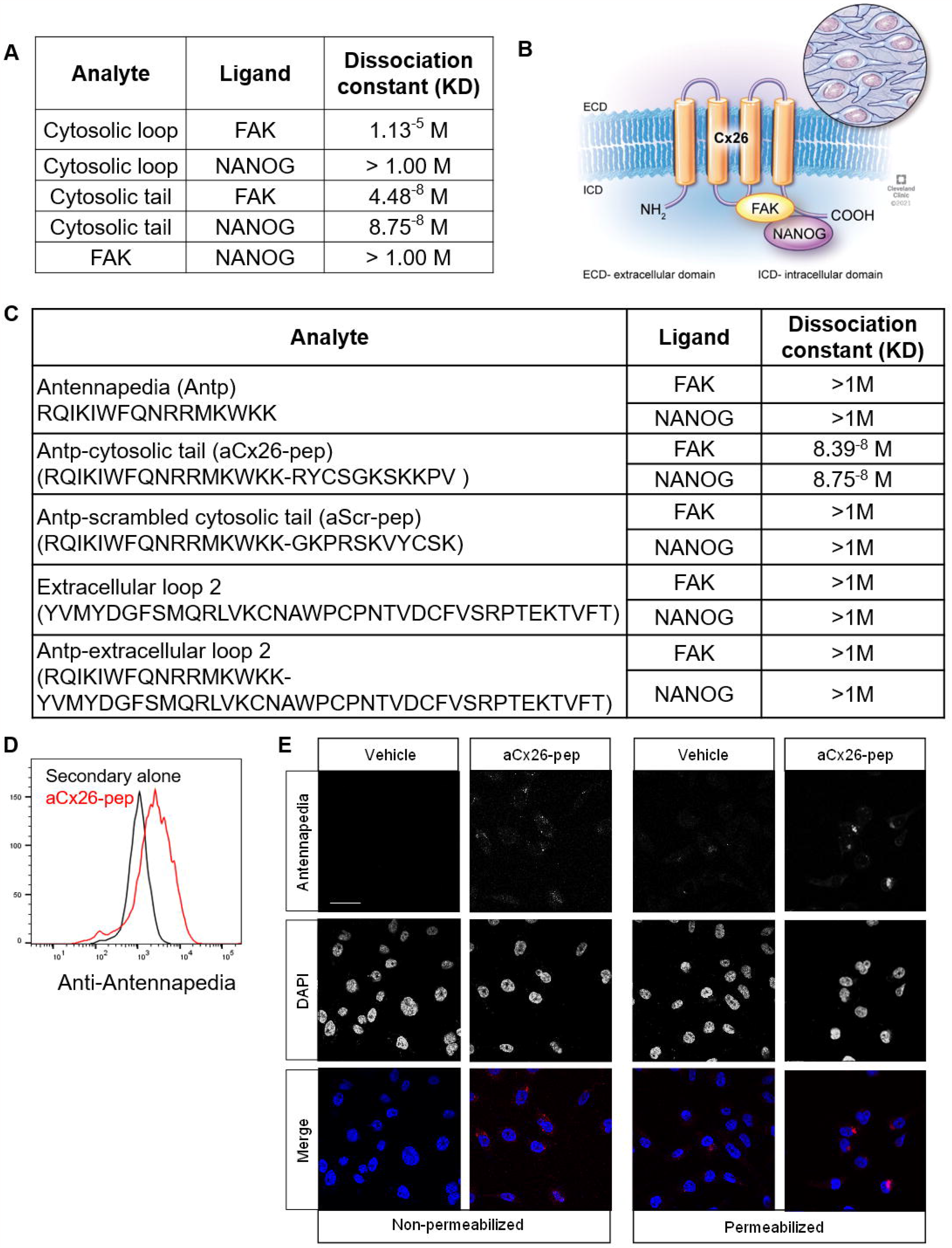
Cx26 cytosolic tail peptide binds both FAK and NANOG and penetrates the cell membrane upon addition of an antennapedia (Antp) leader sequence. A) Table denoting dissociation constants for interactions between Cx26 and NANOG and FAK. B) Cartoon depiction of complex based on SPR analysis. C) Summary table of FAK and NANOG dissociation constants with Antp alone, tagged cytosolic tail, or tagged cleaved tail peptides by SPR analysis. D) Intracellular flow cytometry staining and E) immunofluorescence staining for Antp following 10 μM aCx26-pep treatment for 16 h, demonstrating the ability of the peptide to cross the plasma membrane. Scale bar, 40 μm.

Based on the observation that the Cx26 cytosolic tail is essential for interaction with NANOG and FAK, we sought to understand the effect of disrupting the complex using a Cx26 cytosolic tail-derived cell-penetrating peptide. We tagged a peptide containing the Cx26 cytosolic tail sequence with an antennapedia (Antp) (RQIKIWFQNRRMKWKK) leader sequence (termed hereafter aCx26-pep). Antp is a well-studied leader sequence used to allow peptides and small molecules to penetrate the plasma membrane^39^. aCT1, a Cx43 C-terminal tail mimetic peptide, impairs the malignant progression of luminal breast cancer cells and enhances the activity of the standard-of-care chemotherapies lapatinib and tamoxifen^40, 41^. We first confirmed that the Antp leader sequence did not significantly alter the ability of aCx26-pep to bind to either FAK or NANOG by SPR (**Fig. 2C**). To determine the optimal treatment conditions, cells were assessed for the presence of aCx26-pep by flow cytometry (**Fig. 2D**) and intracellular immunofluorescence (IF) staining (**Fig. 2E**) using an anti-Antp antibody. Optimal treatment conditions were determined to be 10 μM peptide for a minimum of 16 h.

### Self-renewal but not proliferation of TNBC cells is attenuated by aCx26-pep treatment

To assess the functional impact of treatment, we evaluated cell proliferation and self-renewal in TNBC cells treated with aCx26-pep in vitro. MDA-MB-231 TNBC cells were treated with increasing doses of vehicle, Antp alone, and aCx26-pep, and cell proliferation was measured after 72 h of treatment. Interestingly, cell proliferation was not significantly affected by treatment with aCx26-pep (**Fig. 3A, B**). To assess changes in self-renewal, tumorsphere formation was interrogated in three TNBC cell lines (MDA-MB-231, MDA-MB-468 and HCC70) and one luminal breast cancer cell line (MCF7) treated with vehicle, Antp, Antp-tagged scrambled aCx26-pep (aScr-pep), and aCx26-pep. Tumorsphere-initiation frequency was assessed using an extreme limiting-dilution assay (ELDA) and subsequently analyzed using ELDA analysis software^37^. TNBC cells treated with aCx26-pep had significantly decreased tumorsphere frequency compared to those treated with aScr-pep, whereas the luminal breast cancer cell line showed no significant difference between aScr-pep and aCx26-pep (**Fig. 3C-F, Supplemental Table S2, Supplemental Fig. S4**). A shortened region of the Cx26 cytosolic tail tagged with Antp, aCx26-pep 216-220, showed similar effects to aCx26-pep (**Supplemental Fig. S5**). We then evaluated the effect of aCx26-pep treatment on HCC70 CSCs and non-CSCs enriched using our NANOG-GFP reporter. aCx26-pep treatment significantly reduced tumorsphere frequency in both NANOG-GFP^hi^ CSCs and NANOG-GFP^lo^ non-CSCs (**Supplemental Fig. S6**). Together, these data show that disrupting the Cx26/NANOG/FAK complex using aCx26-pep decreases the self-renewal capacity of TNBC.

**Figure 3.**
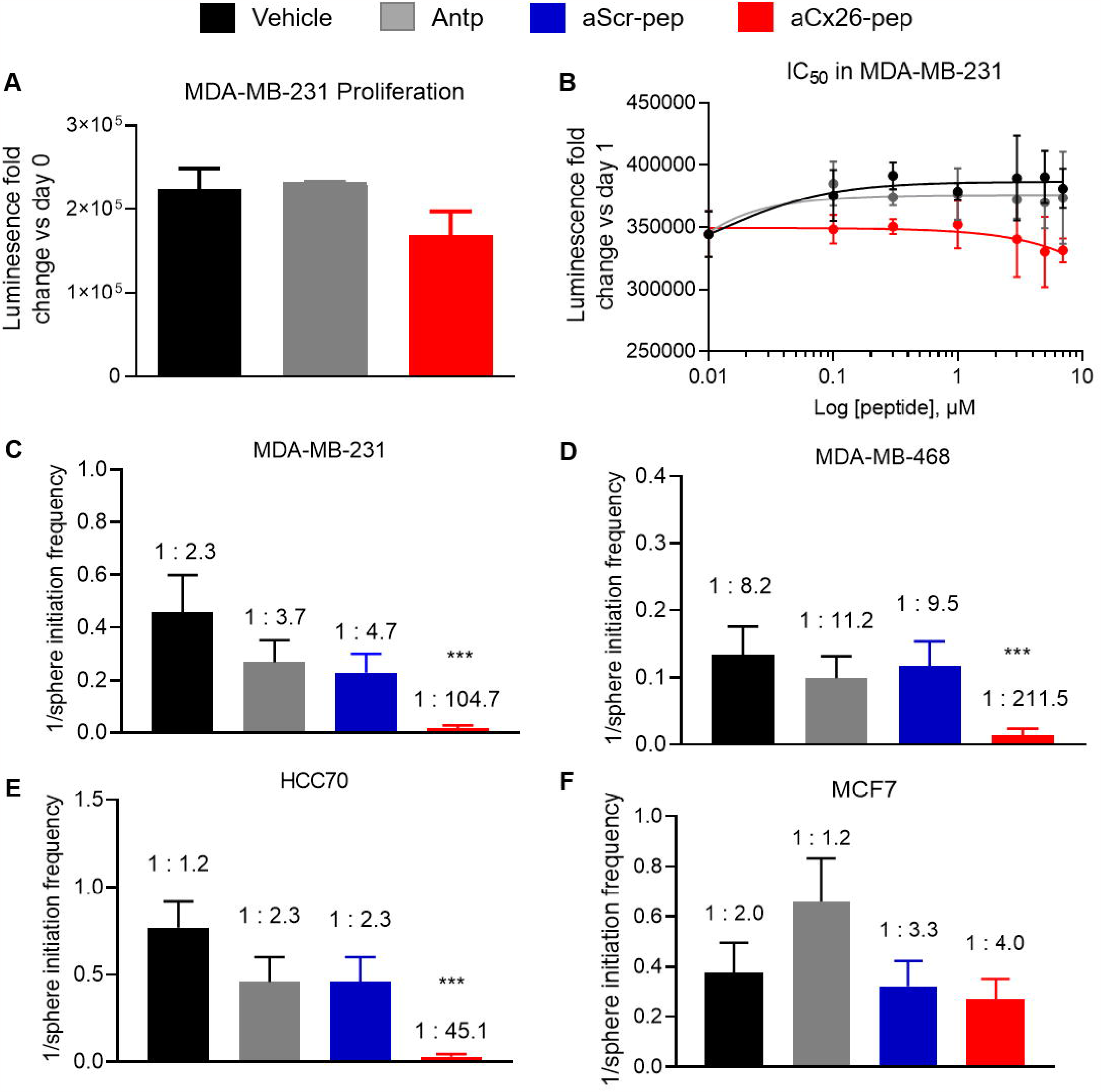
TNBC self-renewal but not cell proliferation is attenuated following aCx26-pep treatment. Cell proliferation measured by CellTiter Glo is not significantly affected by aCx26-pep treatment compared to controls over time (A) or by concentration of peptide (B). Tumorsphere initiation frequency is decreased in MDA-MB-231 C), MDA-MB-468 D), and HCC70 E) TNBC cells treated with aCx26-pep compared to vehicle, Antp alone, or aScr-pep controls but is not affected in MCF7 luminal breast cancer cells (F). *** p<0.001 by Chi-squared test compared to aScr-pep.

### aCx26-pep treatment alters FAK and NANOG levels and NANOG function

To evaluate the effect of aCx26-pep on members of the Cx26/FAK/Nanog complex, we first measured the effect of the peptide on pluripotency transcription factors. While we did not observe a change in *Sox2* levels (*not shown*), there was a significant decrease in Oct4 mRNA levels in both MDA-MB-231 and HCC70 cells treated with aCx26-pep compared to aScr-pep (**Fig. 4A**). We next interrogated whether peptide treatment affected the levels and subcellular localization of NANOG and FAK. After 48 h treatment with aCx26-pep, HCC70 cells were subjected to subcellular fractionation and immunoblotting for FAK and NANOG (**Fig. 4B**). We observed dramatic decreases in both FAK and NANOG levels in the nuclear fraction of TNBC cells, suggesting that treatment with aCx26-pep may disrupt the function of these proteins. This recapitulates the destabilization of NANOG we previously observed after knockdown of Cx26^34^. To further understand the downstream signaling changes following treatment of TNBC cells with aCx26-pep, we performed a targeted qPCR assessment of NANOG-regulated genes. Following 7 days of treatment with vehicle, aScr-pep, or aCx26-pep, expression of NANOG target genes including *FABP5, POSTN, NRP2, PIN1*, and *TAGLN* was significantly decreased in MDA-MB-231 TNBC cells treated with aCx26-pep compared to those treated with vehicle or aScr-pep (**Fig. 4C**). Additionally, expression of epithelial cell markers such as cytokeratin 18 and 19 (*KRT18* and *KRT19*) was significantly increased in the aCx26-pep treatment group (**Fig. 4D**). In contrast, treatment of MCF7 luminal breast cancer cells did not significantly change NANOG target gene or cytokeratin gene expression (**Fig. 4D-F**). Together, these results indicate decreased levels of FAK and NANOG, reduced transcriptional activity of NANOG, and an increase in epithelial cell signature after treatment with aCx26-pep.

**Figure 4.**
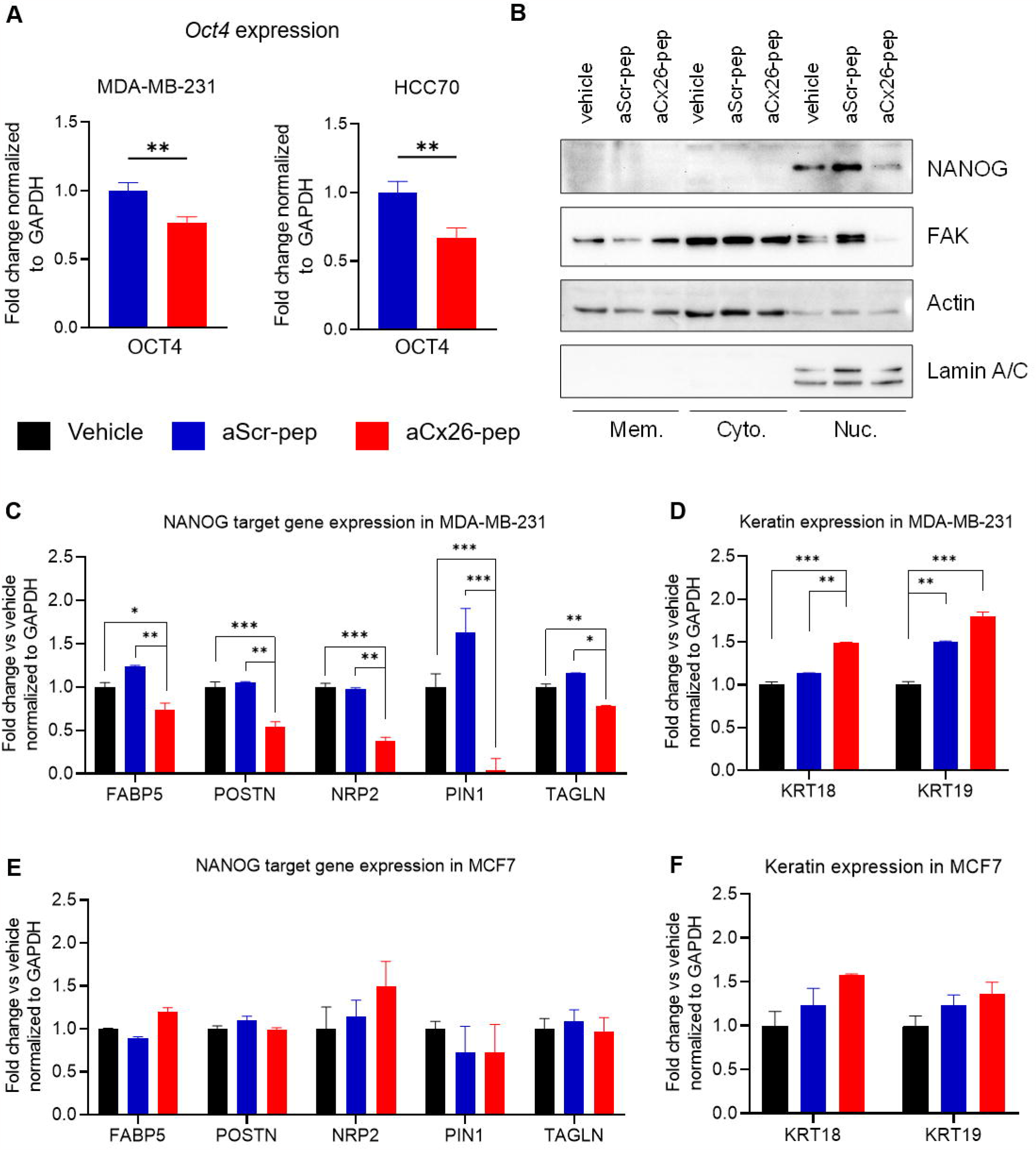
aCx26-pep treatment alters FAK and NANOG levels and NANOG function. A) Following a 7 day treatment with 10 μM aCx26-pep, MDA-MB-231 and HCC70 cells had decreased expression of the pluripotency transcription factor *Oct4*. B) Treatment with aCx26-pep for 48 h decreased the amount of FAK and NANOG in the nuclear fraction. Actin and Lamin A/C were used as loading controls. Mem., membrane. Cyto, cytoplasmic. Nuc., nuclear. C-F) After 7 days of treatment, genes regulated by NANOG by qPCR were decreased with aCx26-pep compared to cells treated with either cytosolic tail peptide alone, aScr-pep or vehicle alone (C), while the expression of keratin genes was increased (D). There were no significant differences in gene expression between the vehicle, cytosolic tail, or aScr-pep groups. E-F) No changes were not observed in NANOG target or keratin gene expression in MCF7 luminal breast cancer cells. n=3 independent experimental replicates, each performed in technical triplicate. * p<0.05, ** p<0.01, *** p<0.001 by one-way ANOVA.

### In vivo TNBC tumor progression is attenuated by aCx26-pep

To determine the effect of aCx26-pep on tumor formation *in vivo*, NSG mice were subcutaneously injected into the flank with TNBC MDA-MB-231 luciferase-expressing cells and randomized into treatment groups once tumors became palpable. Mice were intra-tumorally injected every other day for 20 days with either vehicle, aScr-pep, or aCx26-pep and monitored by weekly bioluminescence imaging (**Fig. 5A**). Mice treated with aCx26-pep exhibited significantly reduced luciferase signal (tumor burden) over time when compared to those treated with aScr-pep or vehicle controls (**Fig. 5B, C**).

**Figure 5.**
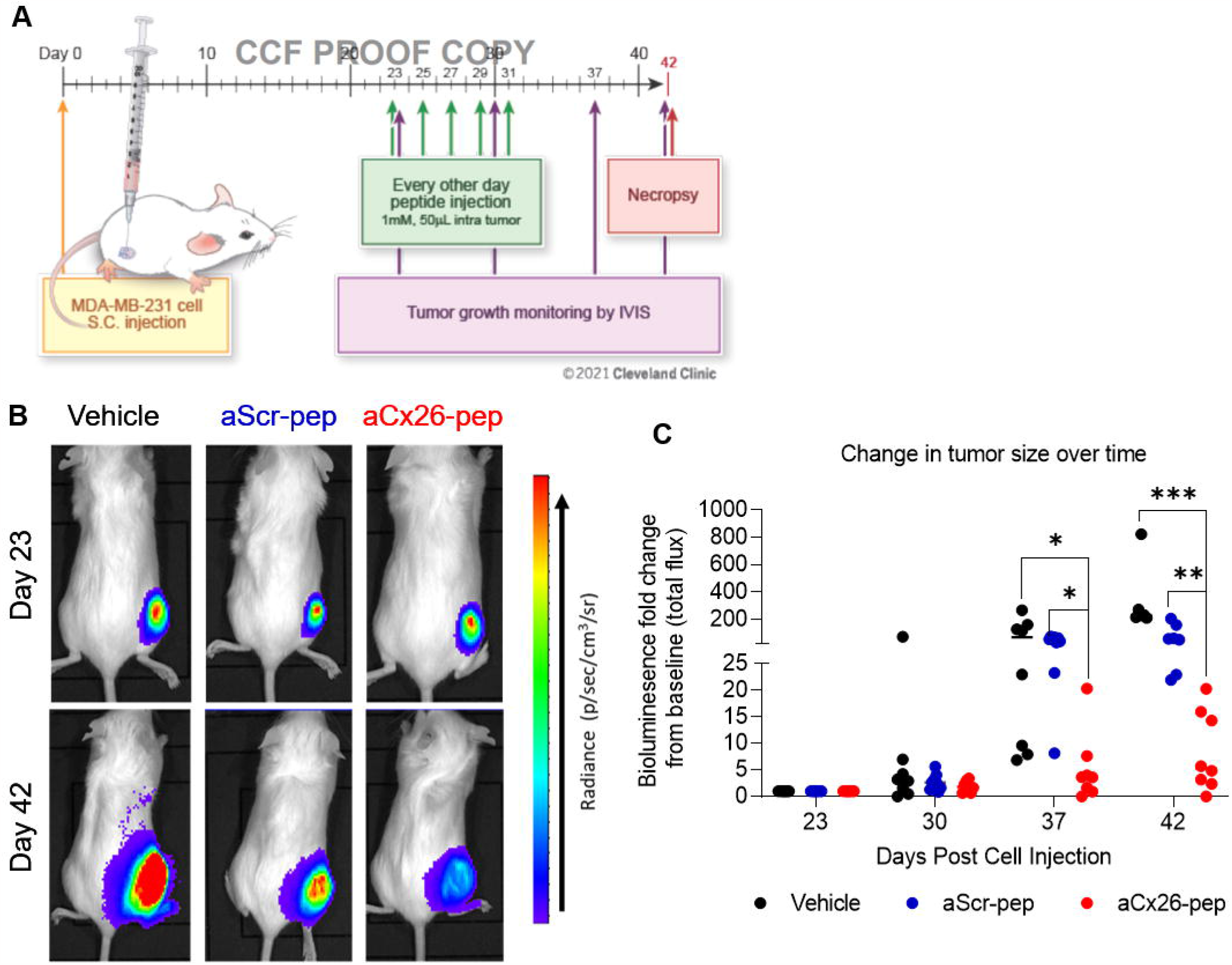
Tumor growth is attenuated upon aCx26-pep treatment in vivo. A) Following palpability of tumors in the subcutaneous right flank generated by MDA-MB-231 cell injection, mice were intratumorally treated with 50 μl of either 0.1 mM peptide or vehicle (1% DMSO) every other day for 20 days and monitored by weekly IVIS imaging. B) Representative IVIS images of treated mice at Day 0 and Day 42. C) Tumors treated with aCx26-pep exhibited attenuated tumor growth over time compared to control groups. n=8 mice per group; * p<0.05, ** p<0.01, *** p<0.001 by two-way ANOVA.

We then performed histological assessment of the resulting tumors. Upon H&E staining of isolated tumors, we observed that the percent tumor viability was significantly decreased in aCx26-pep-treated mice compared to vehicle- or aScr-pep-treated mice (**Fig. 6A**). Immunofluorescence staining of the endpoint tumors showed that Ki67 (**Fig. 6B**), a marker for proliferation, and vimentin (**Fig. 6C**), a marker for epithelial-mesenchymal transition (EMT), were decreased, while cleaved caspase 3 was increased (**Fig. 6D**), in the aCx26-pep-treated groups, suggesting that peptide treatment led to a more epithelial cell phenotype with increased cell death and diminished proliferation. Taken together, these data support the efficacy of aCx26-pep for targeting TNBC CSCs.

**Figure 6.**
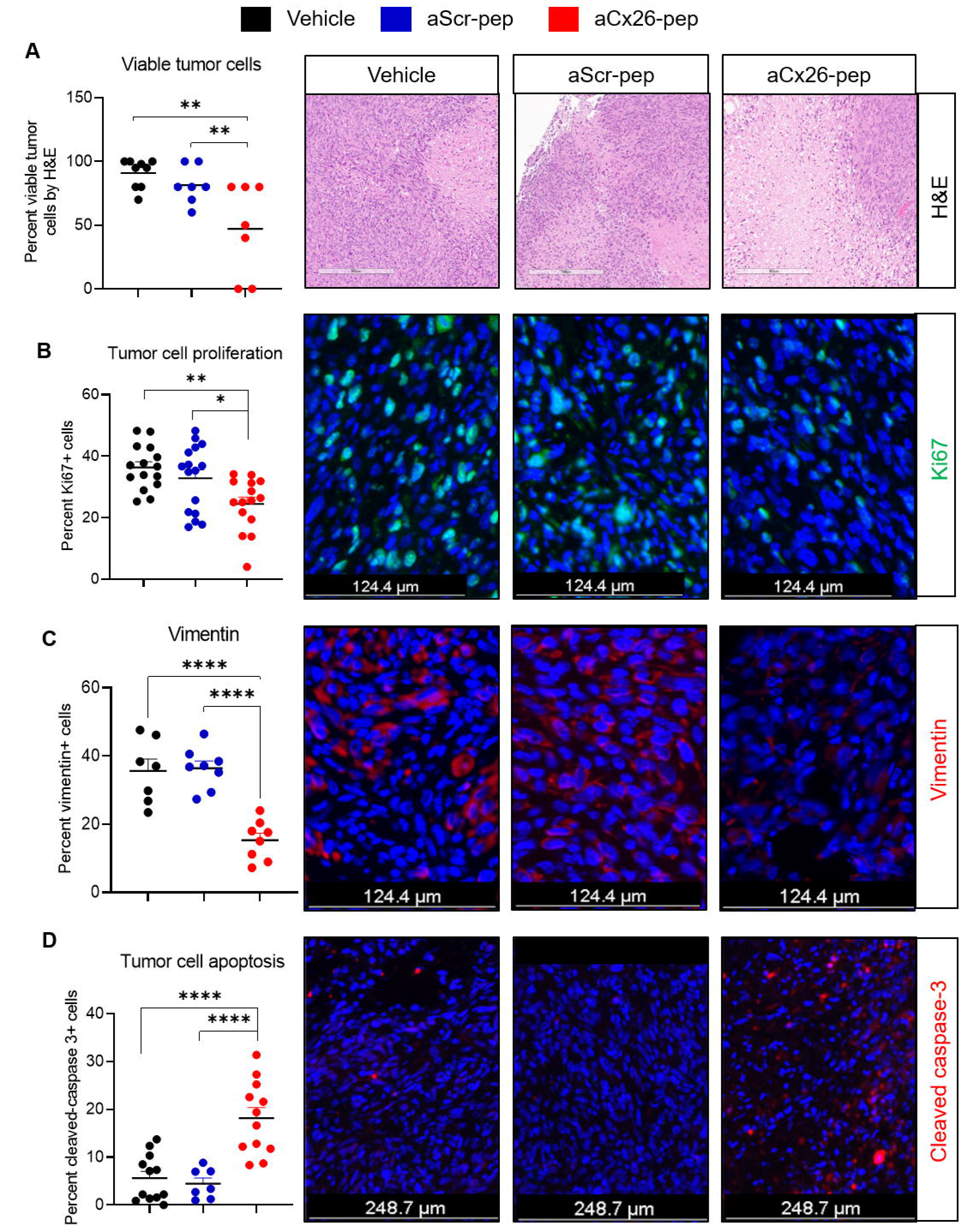
aCx26-pep-treated tumors show decreased tumor proliferation and cell viability and increased cell death. At endpoint, aCx26-pep-treated tumors exhibited diminished viability by H&E scoring (A), decreased Ki67 staining (B), decreased vimentin expression (C), and increased cleaved caspase 3 levels (D) compared to tumors treated with either aScr-pep or vehicle. H&E: 7-8 mice per group, IF staining: 3-4 mice per group, 3 images per mouse. Scales bars are as denoted in each fluorescence image; for (A), scale bar = 302 μm. *p<0.05, **p<0.01, ***p<0.001, ****p<0.0001 by one-way ANOVA.

## Discussion

Overall, our findings provide proof-of-concept data that a peptide strategy can be employed to reduce CSC self-renewal and tumor growth. We leveraged previous observations of the Cx26/NANOG/FAK complex in TNBC to develop a pre-clinical peptide-based targeting strategy to attenuate CSC function. Our studies focused solely on CSCs and tumor growth, key cancer phenotypes that likely involve FAK and NANOG. However, given the number of downstream signaling networks associated with FAK and NANOG, there are likely to be other cellular programs that may also be driven by their association with Cx26, including adhesion, migration, and invasion. An additional unanswered question is the possibility of differential functions of free and Cx26-complex-bound NANOG and FAK.

Immunohistochemical analysis complemented our breast cancer cell line data, with the majority of TNBC specimens having cytoplasmic expression of Cx26 and a subset exhibiting a similar staining pattern of FAK and NANOG. In contrast, the majority of HR+/HER2-carcinomas showed membranous expression of Cx26 and cytoplasmic localization of FAK and NANOG. The distinct cellular localization of Cx26 expression in TNBC and HR+/HER2-tumors further supports the hypothesis that therapeutic targeting of Cx26 would benefit this aggressive tumor subtype. Although the immunohistochemical TNBC data are not as compelling as those seen in the TNBC cell lines, FFPE immunohistochemistry may suffer from antibody affinity, fixation issues, and antigen retrieval. Clinical outcomes data related to a larger cohort defined with Cx26, FAK and NANOG will be helpful in further elucidating this complex and its role.

Our findings identify a strategy to target CSCs and the NANOG-associated signaling network by blocking intracellular Cx26 interactions via a cell-penetrating peptide approach. A challenge in targeting connexin family members remains specificity, as there is a high degree of structural homology among connexin family members, apart from the cytoplasmic tail that provides a binding surface for adaptor proteins to initiate signaling programs. In previous work, the cytoplasmic tail of Cx43 was found to associate with zona occludens-1 (ZO-1) to exclude assembled connexins from the gap junction plaque; disruption of the interaction with ZO-1 resulted in an increase in gap junction intracellular communication^41^. A peptide mimetic of this region of the Cx43 tail tagged with antennapedia has been successfully leveraged to enhance diabetic would healing and is in advanced stages of clinical testing^42^, and a TAT fusion of a region of Cx43 that binds to SRC has been shown to decrease glioblastoma CSC phenotypes^43-47^. These findings provided the first example that peptides can be used to modulate intracellular connexin function for a clinically relevant application. Our findings provide the first evidence of the utility of this application for TNBC targeting and potential downstream signaling modulation. While these peptide-based approaches show promise, there remain barriers to implementation, including delivery and stability. While we primarily tested aCx26-pep, which contains the full 11 amino acid cytosolic tail sequence, we observed similar effects with the first five amino acids of the cytosolic tail, RYCSG, which corresponds to amino acids 216-220. Further study is required to determine the optimal peptide sequence for both efficacy and stability. Differences in concentrations of peptide available, as well as the presence of the tumor microenvironment, may have been responsible for slight differences between peptide effects observed in vivo and in vitro, such as tumor cell viability. An additional question is the effect of our targeting peptide on normal stem cell function. Based on our observation that aCx26-pep does not affect luminal breast cancer cells such as MCF7, which also express FAK and NANOG, we would anticipate limited toxicity to normal stem cells. Future studies will focus on addressing these key limitations to facilitate clinical translation.

Targeting CSCs remains of interest in many cancers, including TNBC, but many late-stage trials are focused on canonical developmental programs (including Sonic hedgehog, Wnt, and Notch). Next-generation strategies are more likely to be successful if they can leverage CSC-specific signaling networks. CSC targeting represents a new cancer therapeutic development pipeline that is complementary to oncogene addiction targeting and tumor suppressor re-expression. Connexins and their main functions of cell-cell communication, hemichannel communication, and initiation of signaling have been largely understudied in the context of cancer and may represent a new therapeutic target. These findings highlight a selective strategy using a peptide approach to target TNBC, leading to new therapeutic options for the unmet need in this patient population.

## Acknowledgements

The authors would like to thank members of the Lathia and Reizes laboratories for insightful comments during implementation of these studies. We appreciate the technical contributions of Chad Braley; members of the Lerner Genomics Shared Lab Resources, specifically Dr. Dean Horton; and the Imaging Core, namely Dr. Judith Drazba. The research was supported by the Department of Defense Grant W81XWH1910503, NIH R21 CA191263, and VeloSano Bike to Cure. Work in the Lathia laboratory is also supported by NIH R35 NS127083 and P01 CA245705, the Lerner Research Institute, and the Case Comprehensive Cancer Center. Work in the Reizes laboratory is also supported by Cleveland Clinic Center of Excellence, VeloSano Bike to Cure and The Laura J. Fogarty Endowed Chair for Uterine Cancer Research.

