## Supplemental figures for "Targeting the Cx26/NANOG/Focal Adhesion Kinase Complex via Cell-Penetrating Peptides in Triple-Negative Breast Cancers": Supplemental information.pdf

**Supplemental Figure S1**. Predicted model of Cx26 in complex with NANOG and FAK.

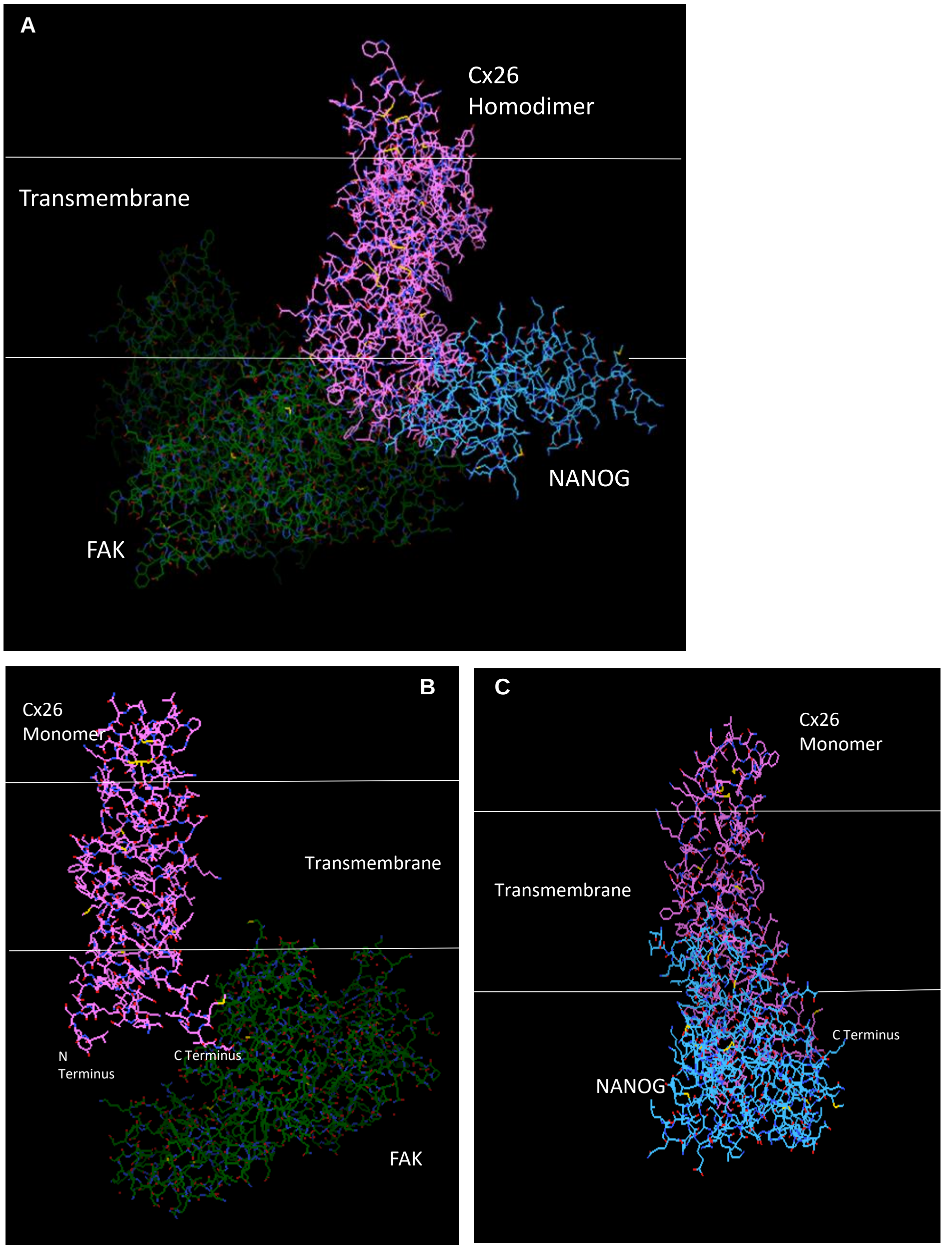

**Supplemental Figure S2.** SPR sensograms demonstrating binding of Cx26 peptides to FAK or NANOG proteins.

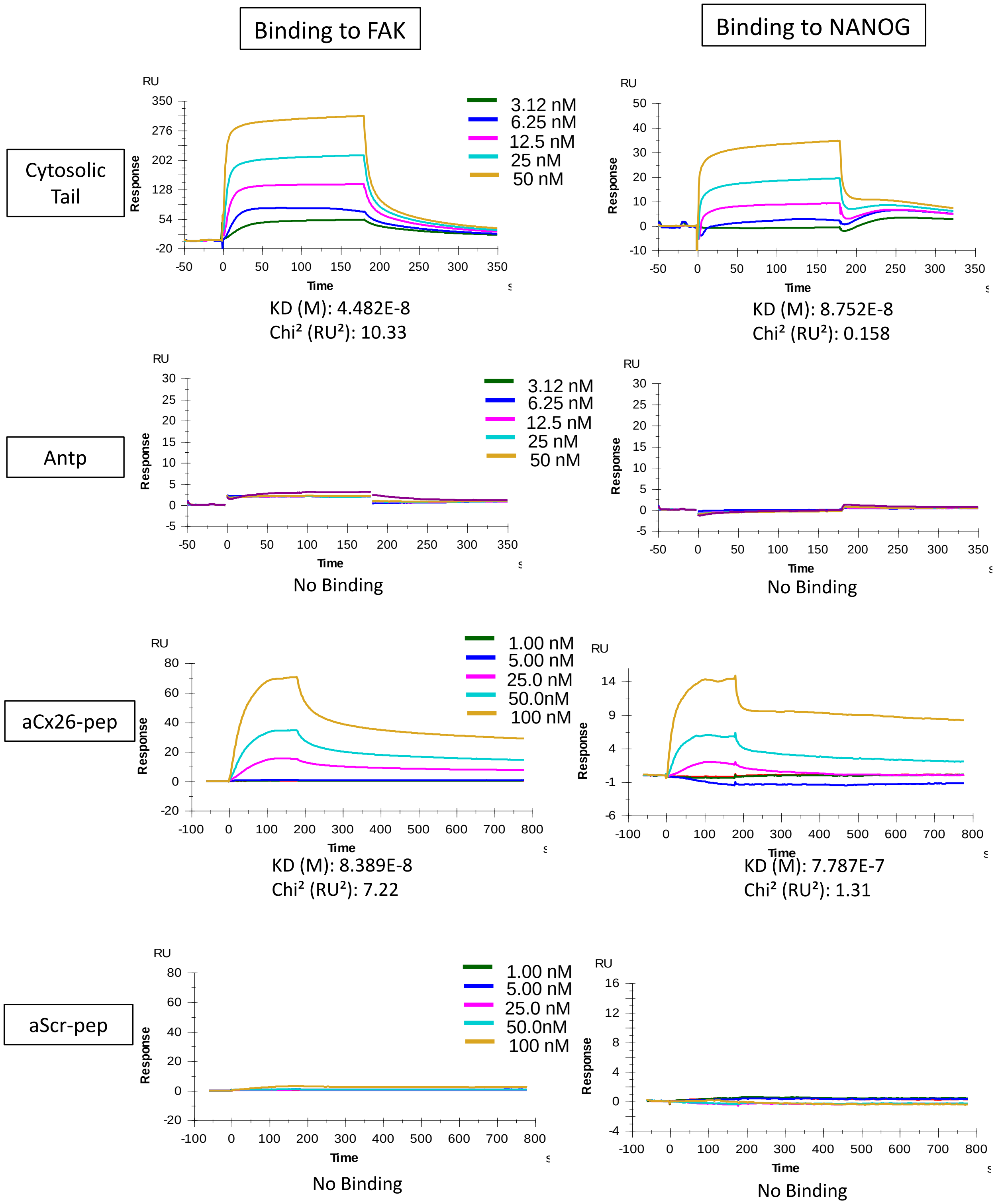

**Supplemental Figure S3.** SPR sensograms demonstrating no binding of Cx26 extracellular loop peptides to FAK or NANOG proteins.

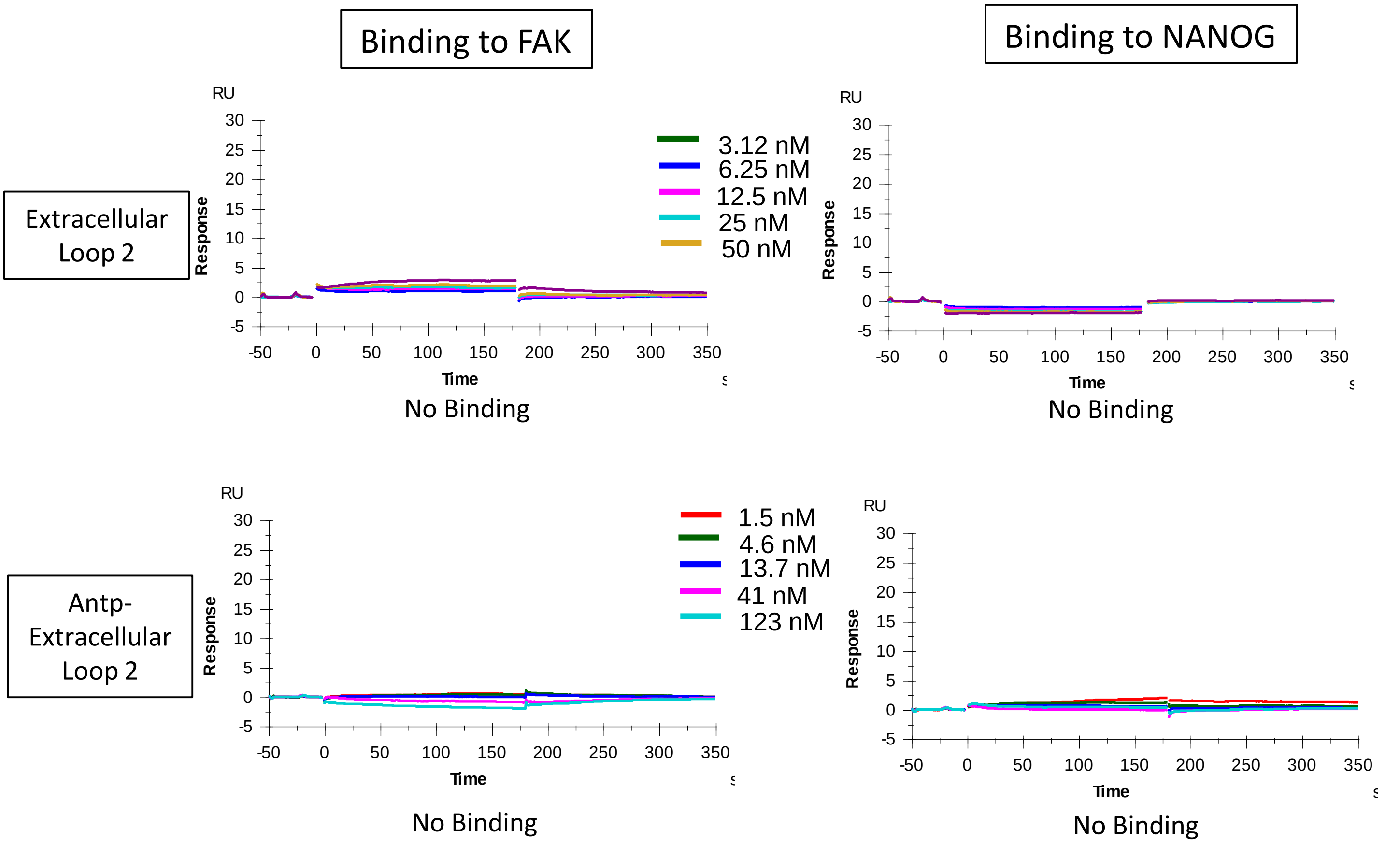

**Supplemental Figure S4.** Representative images from the sphere-forming assays shown in Fig. 3C (MDA-MB-231). Scale bar, 400 μm.

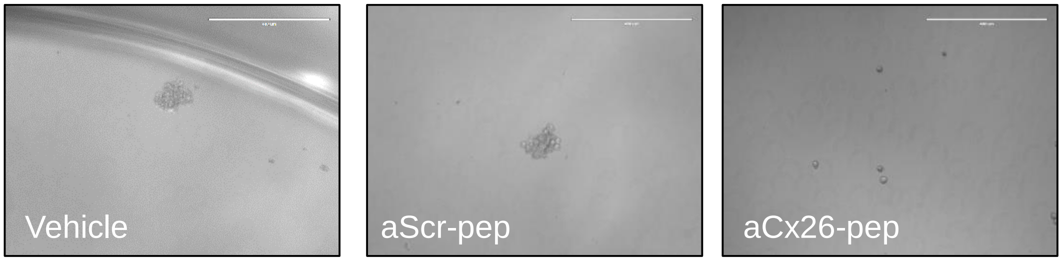

**Supplemental Figure S5.** The shortened aCx26-pep 216-220 attenuates sphere-forming frequency similarly to the full-length aCx26-pep. *** p<0.001 by Chi-squared test compared to aScr-pep. Shown is one replicate representative of three independent experiments.

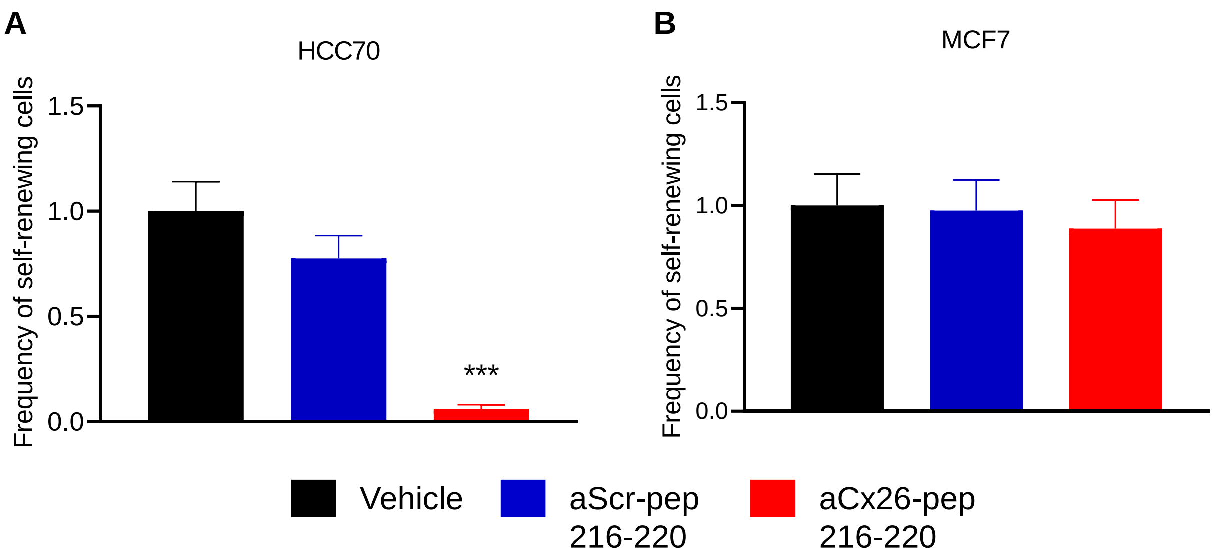

**Supplemental Figure S6.** Self-renewal of both HCC70 NANOG-GFP$^{hi}$ and NANOG-GFP$^{lo}$ cells is reduced by treatment with aCx26-pep. Shown is one representative replicate from three independent experiments. *** $p<0.001$ by Chi-squared test compared to aScr-pep.

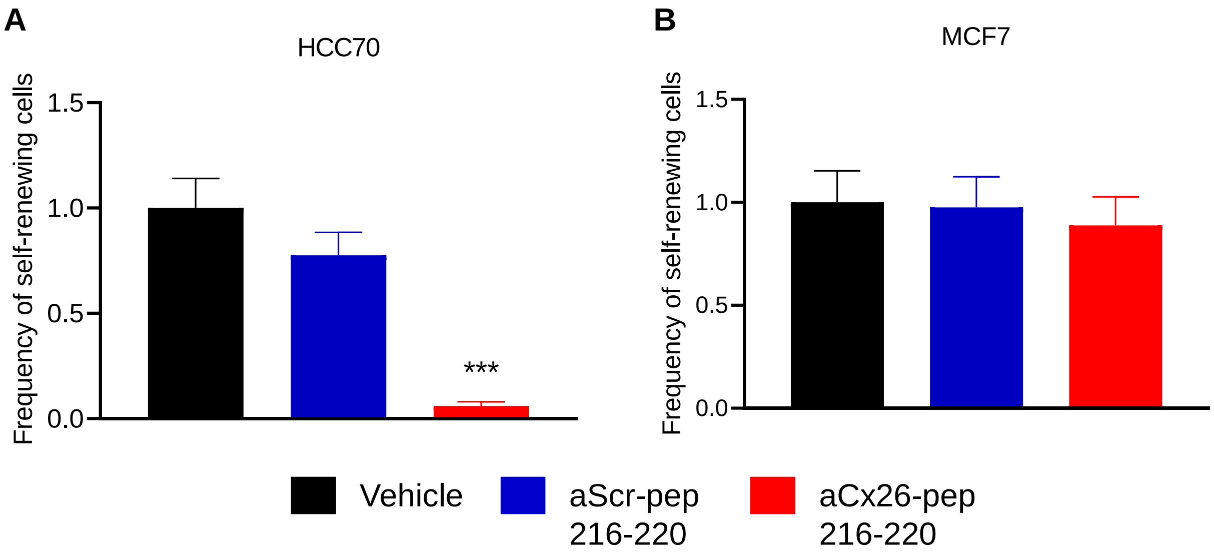

**Supplemental Tables.**

**Supplemental Table S1.** Dissociation constants derived from SPR showing that FAK and NANOG specifically to amino acids 216-220 of the Cx26 cytosolic tail.

| **Name** | **Sequence** | **NANOG** | **FAK** |
|---|---|---|---|
| Wildtype | RYCSGKSKKPV | $1.80^{-7}$ M | $1.49^{-5}$ M |
| xR216 | XYCSGKSKKPV | $5.38^{-7}$ M | $2.0^{-5}$ M |
| xRY | XXCSGKSKKPV | $8.76^{-6}$ M | $1.94^{-6}$ M |
| 221-226 | XXXXXKSKKPV | > 1.00 M | > 1.00 M |
| 216-220 | RYCSGXXXXXX | $6.40^{-9}$ M | $4.5^{-7}$ M |
| xV226 | RYCSGKSKKPX | $2.10^{-7}$ M | $2.03^{-5}$ M |

**Supplemental Table S2.** Estimates for sphere-forming efficiency and p-values from Fig. 3C-F.

| Cell Line | Peptide Treatment | Confidence intervals for 1/(stem cell frequency) | | | p-value |
|---|---|---|---|---|---|
| | | Lower | Estimate | Upper | |
| MDA-MB-231 | Vehicle | 4.39 | 2.3 | 1.4 | - |
| MDA-MB-231 | Antp | 7.27 | 3.7 | 2.37 | n.s. |
| MDA-MB-231 | aScr-pep | 8.43 | 4.73 | 2.78 | n.s. |
| MDA-MB-231 | aCx26-pep | 427.39 | 104.7 | 25.94 | 4.0e-12 |
| MDA-MB-468 | Vehicle | 14.6 | 8.2 | 4.8 | - |
| MDA-MB-468 | Antp | 20.4 | 11.2 | 6.3 | n.s. |
| MDA-MB-468 | aScr-pep | 17 | 9.46 | 4.7 | n.s. |
| MDA-MB-468 | aCx26-pep | 1514.6 | 211.5 | 29.9 | 6.7e-5 |
| HCC70 | Vehicle | 2.1 | 1.2 | 1.0 | - |
| HCC70 | Antp | 4.4 | 2.3 | 1.4 | n.s. |
| HCC70 | aScr-pep | 4.3 | 2.31 | 1.4 | n.s. |
| HCC70 | aCx26-pep | 118.1 | 45.1 | 17.5 | 6.2e-13 |
| MCF7 | Vehicle | 3.9 | 2.0 | 1.3 | - |
| MCF7 | Antp | 2.1 | 1.2 | 1.0 | n.s. |
| MCF7 | aScr-pep | 6.12 | 3.35 | 1.97 | n.s. |
| MCF7 | aCx26-pep | 7.27 | 4.0 | 2.3 | n.s. |

**Supplemental Table S3.** qRT-PCR primers used in this study.

| Gene Name | Direction | Sequence (5’-3’) |
|---|---|---|
| FABP5 | Forward | TGA AGG AGC TAG GAG TGG GAA |
| | Reverse | TGC ACC ATC TGT AAA GTT GCA G |
| KRT18 | Forward | GGC ATC CAG AAC GAG AAG GAG |
| | Reverse | ATT GTC CAC AGT ATT TGC GAA GA |
| KRT19 | Forward | AAC GGC GAG CTA CAG GTG A |
| | Reverse | GGA TGG TCG TGT AGT AGT GGC |
| NRP2 | Forward | GCT GGC TAT ATC ACC TCT CCC |
| | Reverse | TCT CGA TTT CAA AGT GAG GGT TG |
| PIN1 | Forward | TGC CAC CGT CAC ACA GTA TT |
| | Reverse | CTG ACC AGG CCT TCT CTT TG |
| POSTN | Forward | CTC ATA GTC GTA TCA GGG GTC G |
| | Reverse | ACA CAG TCG TTT TCT GTC CAC |
| TAGLN | Forward | AGT GCA GTC CAA AAT CGA GAA G |
| | Reverse | CTT GCT CAG AAT CAC GCC AT |
| OCT4 | Forward | TGA GTC AGT GAA CAG GGA ATG |
| | Reverse | AAT CTC CCC TTT CCA TTC GG |
